# Novel insights into the biosynthesis and diversity of leaderless multipeptide bacteriocins

**DOI:** 10.64898/2026.09.04.749385

**Authors:** Thomas F. Oftedal, Simen Hermansen, Even A. Haslund, Amar A. Telke, Tage Thorstensen, Kirill V. Ovchinnikov, Dzung B. Diep, Morten Kjos

## Abstract

Garvicin KS (GarKS) is a three-peptide, leaderless, broad-spectrum bacteriocin that is active against a wide range of Gram-positive bacteria, including several foodborne pathogens and antibiotic-resistant strains. This bacteriocin is considered a candidate for application in food preservation and medical treatments; however, key knowledge about the producer strain, regulation of production, and the mechanism of action is still lacking for leaderless, multipeptide bacteriocins, including GarKS. A hybrid sequencing strategy was used to obtain a high-quality closed genome assembly of the native GarKS producer *Lactococcus garvieae* KS1546, which showed that the bacteriocin was encoded on a 50 kb plasmid (pKS50). Comparative analysis with updated sequence databases indicates that the producer strain should be reclassified as *Lactococcus petauri*. The roles of the biosynthetic genes, putatively encoding a transcriptional regulator (*gakR*) and immunity protein (*gakI*), were examined using heterologous expression. Removal of *gakR* resulted in a 6-fold decrease in GarKS production, and expression of *gakI* caused a 250-fold decrease in susceptibility to GarKS, with varying degrees of cross-immunity to other multipeptide bacteriocins. Isolation and whole-genome sequencing of spontaneous GarKS mutants in *L. lactis* showed that resistance levels are low, but that *ythA*, a PspC-domain-containing protein involved in a phage stress response pathway is involved in the GarKS susceptible phenotype. To decipher conserved features involved in production of these bacteriocins, we finally used genome mining to identify 11 candidates for new multipeptide bacteriocins; four of them were obtained synthetically and confirmed to be bioactive peptides inhibiting important pathogens including *Listeria monocytogenes* and enterococci.

## Introduction

Bacteriocins are ribosomally synthesized antimicrobial peptides or proteins produced by many Gram-negative and Gram-positive bacteria to gain a competitive advantage for nutrients in their environment ^1^. The greatest diversity of bacteriocins originates from Gram-positive bacteria, with lactic acid bacteria being frequent producers ^2,3^. These molecules are typically small, cationic, and most often thought to exert their antimicrobial activity by disrupting the membrane integrity of target cells, causing leakage of cellular contents and ultimately leading to cell death ^4^. While many bacteriocins have a narrow spectrum of activity, often targeting only closely related species or genera, some exhibit broader antimicrobial spectra ^5–7^. Their targets can include problematic foodborne bacteria such as *Listeria monocytogenes* and *Bacillus cereus*, as well as pathogens like vancomycin-resistant enterococci (VRE) and methicillin-resistant *Staphylococcus aureus* (MRSA) ^8^. Due to these properties, bacteriocins hold significant promise as food biopreservatives and potential therapeutic agents ^9^.

Bacteriocins exhibit considerable diversity in their structure, biosynthesis, and mechanisms of action. Some are post-translationally modified and often referred to as class I bacteriocins. These include the lantibiotics, which contain polycyclic thioether amino acids like lanthionine or methyllanthionine ^10^. Class II bacteriocins consist of unmodified and structurally simpler peptides, which are further divided into subclasses, including, the class IIa pediocin-like bacteriocins ^11^ and class IId single-peptide, non-pediocin-like bacteriocins, such as lactococcin A and garvieacin Q ^12–15^. While many bacteriocins, including class IIa and IId, are active as single peptides, others require multiple different peptides for optimal activity ^16^. Bacteriocins that rely on two different peptides are grouped in subclass IIb, also known as two-peptide bacteriocins. Examples of these include the bacteriocins lactococcin G, enterocin DD14 and enterocin L50 ^17–19^. Subclass IIc consists of the so-called leaderless bacteriocins, where the translated peptide is identical to the active secreted peptide, most of which are single peptide bacteriocins ^4,20^. Within subclass IIc, there is however a group of bacteriocins requiring three or four distinct peptides for optimal activity. This group, which is studied in this work, is collectively referred to as the multipeptide bacteriocin family ^7^. Members of this family are the three-peptide bacteriocins garvicin KS (GarKS), and cereucins V, X, and H (CerV, X, H) ^7^, as well as the four-peptide aureocin A70 (AurA70) ^21^.

The genes involved in bacteriocin production, immunity, and export are typically organized in clusters, often comprising multiple operons ^22^. Bacteriocin production may be regulated by two-component regulatory systems composed of a histidine kinase and a response regulator, enabling quorum-sensing control of bacteriocin biosynthesis ^3,23–26^. In some cases, bacteriocin gene clusters encode DNA-binding proteins unrelated to two-component systems ^7,27,28^. These proteins are less characterized, but often bind to inverted repeats within or near promoter regions to regulate transcription. Examples include CylR1 and CylR2 involved in regulating the production of cytolysin, and AurR, which is thought to downregulate production of aureocin A70 ^29–31^.

Most bacteriocins are synthesized as precursors containing a leader peptide, typically at the N-terminus, which is important for recognition by modification enzymes and/or export proteins ^32–34^. Leader peptides are thought to keep bacteriocins in an inactive form prior to export out of the cell. This peptide export often occurs concomitant with the removal of the leader peptide. For leaderless bacteriocins (class IIc), the mechanism of intracellular immunity and recognition by exporters is poorly understood ^20,35^.

To avoid self-killing by their own bacteriocins after secretion, producer strains possess immunity systems, involving expression of dedicated immunity proteins ^35–37^. While such proteins are identified for most bacteriocins, such as AurI for Aureocin A70 ^38^, the mechanism of immunity is not understood for most class II bacteriocins, including the multipeptide bacteriocins. Among class II bacteriocins, the best characterized immunity mechanism is for bacteriocins of class IIa and some of class IId that target the mannose phosphotransferase system (Man-PTS) to kill target cells. The immunity proteins for these bacteriocins bind to subunits of the (Man-PTS) to provide immunity ^37,39–43^.

We previously isolated a *Lactococcus garvieae* strain (KS1546) from raw milk that produces the multipeptide, leaderless bacteriocin called garvicin KS ^7^. GarKS consists of three distinct peptides, each comprising 30–34 amino acids ^7^. GarKS exhibits a broad antimicrobial spectrum, inhibiting not only species within the genus *Lactococcus* but also more distantly related genera including food spoilage bacteria and foodborne pathogens such as *Lactobacillus*, *Streptococcus*, *Listeria*, *Enterococcus*, *Staphylococcus* and *Bacillus* ^7,44–47^. The mechanism of action of GarKS is generally assumed to involve interference with the bacterial membrane, but this has not yet been directly demonstrated experimentally. In *L. monocytogenes*, the phage shock protein response was shown to modulate the susceptibility to GarKS, but the rates and level of resistance towards *L. monocytogenes* were demonstrated to be very low ^48^. Furthermore, high-yield production of GarKS has been achieved by batch fermentation using inexpensive growth medium ^49^. Due to these characteristics, GarKS and the producer strain show strong potential for industrial applications, particularly as a food biopreservative. However, the biosynthesis, immunity, and mode of action of GarKS are not yet fully understood.

To support future applications of GarKS, a comprehensive characterization of GarKS and its producing strain is necessary. In this study, we assembled a complete genome sequence for *L. garvieae* KS1546. Using a heterologous host for expression of different variants of the locus, we experimentally demonstrated which genes are responsible for GarKS immunity and regulation of its biosynthesis. Finally, through genome mining we also identified several new members of the multipeptide bacteriocin family, which enhance our understanding of this unique group of peptides.

## Materials and methods

### Strains and growth conditions

*Lactococcus garvieae* KS1546 ^7,50^ and *L. lactis* IL1403 ^51^ were grown in M17 broth supplemented with 0.5% glucose (GM17) and de Man, Rogosa and Sharpe (MRS) media at 30 °C without shaking. When necessary, erythromycin (Sigma-Aldrich, St. Louis, MO, USA) was added to 200 µg/ml for *E. coli* and 5 µg/ml for lactococci. *L. lactis* harboring pNZ9530 ^52^ was grown with 5 µg/ml erythromycin and chloramphenicol was used at 5 µg/ml for pNZ8037-derived plasmids ^53^.

### Whole-genome sequencing and assembly of *L. garvieae* KS1546

Genomic DNA was isolated from 3 ml overnight culture of *L. garvieae* KS1546 (GM17, 30 °C) using GenElute bacterial genomic DNA kit (Sigma-Aldrich). Sequencing library preparation was performed with Rapid barcoding kit 24 v14 (SQK-RBK114.24, Oxford Nanopore Technologies, Oxford, UK) using 200 ng of isolated genomic DNA. The prepared sequencing library was loaded onto a MinION Mk1B as part of a pool of 9 samples and sequenced until the flow cell (R10.4.1) was depleted. The raw data was basecalled with Dorado (v1.1.1) using the highest quality model (dna_r10.4.1_e8.2_400bps_sup@v5.2.0).

A consensus genome was assembled using Trycycler (v0.5.5) ^54^ with multiple long-read assemblers: Flye (v2.9.6-b1802) ^55^, Miniasm+Minipolish (v0.3-r179) ^56^ and Raven (v1.8.3) ^57^, Canu (v2.2) ^58^, NECAT (v0.0.1) ^59^ and NextDenovo (v2.5.2) ^60^. The generated consensus sequence was then polished with Medaka (v2.1.1) and Polypolish (v0.6.0) using the short Illumina reads to reduce errors. The assembled genome was annotated using Bakta v1.11.0 ^61^, assessed for completeness using CheckM ^62^ and taxonomic assignment was performed with GTDB-Tk ^63^.

### Database search for potential virulence and antibiotic resistance genes

The genome of KS1546 was screened for putative virulence factors by BLASTx against the VFDB full dataset (Set B; protein sequences) (https://www.mgc.ac.cn/cgi-bin/VFs/v5/main.cgi), with hits considered significant at ≥ 80% amino acid identity and ≥ 70% query coverage, following EFSA-recommended thresholds ^64–67^. Potential antibiotic resistance genes were identified using AMRFinderPlus v4.2.7 and ResFinder v4.7.2 (ResFinder database v2.6.0) ^68,69^.

### Database search for new multipeptide bacteriocins

The amino acid sequence of all known multipeptide bacteriocins and biosynthetic genes were used a query in BLAST searches against nucleotide entries (tblastn) in databases available through NCBI (nr/nt, core_nt). For BLAST searches against the sequence read archive (SRA), a list of accession numbers of submissions to SRA matching the search criteria “Bacillota”[Organism] AND (“2014/01/01”[Publication Date] : “2023/12/10”[Publication Date]) was downloaded. Assemblies of the corresponding submissions (accession numbers) were downloaded from the recently published Logan database ^70^ using an automated script and the AWS CLI. Using BLAST+ locally, BLAST databases were constructed of all contigs in chunks of 10 000 sequences each which was then combined using an alias. Sequence accessions with potential hits to known multipeptide bacteriocins or their biosynthetic genes were downloaded from SRA and assembled using Unicycler v5.0 ^71^. Potential candidates were assessed and annotated manually in SnapGene assisted by the conserved domain database and bakta ^61,72^.

### Minimal inhibitory concentration assay

Individual peptides constituting each bacteriocin were chemically synthesized (Pepmic Co., Ltd, China) and solubilized in 0.1% (v/v) trifluoroacetic acid to peptide stock concentrations of 4 mM for four-peptide bacteriocins and 3 mM for three-peptide bacteriocins. Peptides were then mixed in equal volumes to prepare a 1 mM bacteriocin stock solution (1 mM per peptide) which was diluted to 0.1 mM prior to use. For partial combinations, an equal volume of 0.1% (v/v) trifluoroacetic was used for the omitted peptide(s).

To determine the minimal inhibitory concentration (MIC), a serial dilution of the antimicrobial was prepared in brain heart infusion broth (BHI) (VWR) in the wells of a 96-well plate (Sarstedt, Germany). Stationary phase cultures of the indicator strain were diluted 50-fold in the wells of the plate to a total volume of 0.2 ml. The 96-well plate was incubated at 30 °C for 24 hours prior to measurement. Growth was measured by absorbance at 600 nm using a spectrophotometer (SpectroStar Nano, BMG Labtech). The MIC was defined as the lowest concentration of antimicrobial inhibiting growth of the indicator culture by 90% or more (MIC_90_) compared to a control with no added antimicrobial. Values are the means of four independent experiments.

### Cloning of GarKS biosynthetic genes

Genomic DNA isolated from *L. garvieae* KS1546 (described above) was used as a template in PCR reactions. The complete GarKS biosynthetic cluster (A2T) was amplified using the primer pairs A2T_SacI_F and A2T_HindIII_R (Table S1); *gakI* was amplified using gakI_SacI_F and gakI_HindIII_R; gakABC used A2T_SacI_F and gakABC_HindIII_R; the cluster without gakT used A2T_SacI_F and gakΔT_SacI_R (Table S1).

Deletion of *gakR* from the cluster (*gakΔR*) was constructed by overlap extension PCR of two fragments, the first fragment was amplified using A2T_SacI_F and gakΔR_R, the second using A2T_HindIII_R and gakΔRT. The two purified fragments were then annealed and extended in a separate PCR. All PCR reactions used Phusion High-Fidelity DNA polymerase (New England BioLabs) following the recommended protocol and an annealing temperature of 59 °C for all reactions.

Obtained PCR fragments were purified from the reaction mixture, or from gel when necessary, using NucleoSpin Gel and PCR Clean-Up kit (Macherey-Nagel). The plasmid pMG36e was isolated from *E. coli* DH5α using E.Z.N.A. Plasmid DNA Mini Kit I (Omega Bio-Tek). Purified PCR fragments and plasmid (approx. 1 µg) were digested separately in 20 µl reactions using FastDigest SacI and HindIII enzymes (Thermo Fisher Scientific) for 16 h at 16 °C in the supplied FastDigest buffer. Following purification of digested fragments as described previously, linearized plasmid and fragment was ligated using T4 DNA ligase (Thermo Fisher Scientific) at room temperature (approx. 22 °C) for 20 minutes, then heat-inactivated at 65 °C for 10 min. Electrocompetent *L. lactis* IL1403 was prepared as described by Holo et al. (1991), a 50 µl aliquot of cells were mixed with 5 µl of ligation mixture and transformed by electroporation.

### Isolation of GarKS resistant mutants

To select for GarKS resistant cells, 0.1 ml of a stationary-phase culture of *L. lactis* IL1403 was spread directly onto BHI agar plates supplemented with GarKS at 2-, 4-, 8-, 10-, and 12-times the MIC_90_ (corresponding to 20, 40, 80, 100, and 120 nM). Plates were incubated overnight at 30 °C. Colonies that appeared on the plates were isolated, cultured, and genomic DNA was isolated as described previously. Whole-genome sequencing was performed by Novogene (Cambridge, UK) using Illumina sequencing. Reads from mutant isolates were mapped to the reference contigs with Snippy, which was run with default settings to identify variants ^73^.

### Cloning of *ythA* and effect of *ythA* on GarKS susceptibility

The *ythA* gene was amplified from *L. lactis* IL1403 in a routine PCR reaction with Phusion High-Fidelity DNA polymerase (NEB) using the primer pair ythA_XbaI_R and ythA_BamHI_F (annealing temperature of 61 °C). Amplified PCR product was purified using NucleoSpin Gel and PCR Cleap-Up kit (Macherey-Nagel) and 1 µg was subsequently digested with XbaI and BamHI in Tango buffer (Thermo Fisher Scientific) for 1 hour at 37 °C. Plasmid pNZ8037 was isolated from *E. coli* DH5α using E.Z.N.A. Plasmid DNA Mini Kit I (Omega Bio-Tek). The strain was grown at 37 °C with shaking (180 rpm) in LB medium supplemented with 30 µg/ml chloramphenicol. Digested fragments were purified as described above, then ligated using T4 DNA ligase (Thermo Fisher Scientific) at 16 °C) for 16 h. After heat-inactivation of the ligation reaction (65 °C, 10 min), *L. lactis* IL1403 was transformed as described above consecutively, first with the plasmid pNZ9350, then with the ligation mixture.

## Results

### Complete genome of *L. garvieae* KS1546

*L. garvieae* KS1546 is a lactic acid bacterium with industrial potential as a food preservative, either as a probiotic in fermented foods or by the use of its bacteriocin GarKS to control foodborne pathogens and spoilage bacteria. The available genome sequence of the natural GarKS producer *L. garvieae* KS1546 is incomplete. To perform a genomic characterization of the strain, complete whole-genome sequencing and comprehensive bioinformatic analysis was performed. Using a consensus-based long-read assembly combined with short-read polishing allowed us to assemble a complete and accurate genome of *L. garvieae* KS1546 (99.02% completeness and 1.62% contamination by CheckM ^62^). The complete genome, visualized in Figure 1, was found to have a length of 2 208 664 bp and consist of five replicons, including four predicted extrachromosomal plasmids that we named pKS2, pKS26, pKS50 and pKS117. Estimated copy-numbers of the plasmids based on sequencing coverage were 40, 4, 2 and 3 respectively. Genome comparisons against the Genome Taxonomy Database (GTDB) showed the highest average nucleotide identity (ANI) to *Lactococcus petauri* (97.32%, conclusive), compared to 93.1% for *L. garvieae*. A summary of features found in the genome is shown in Table 1.

**Figure 1.**
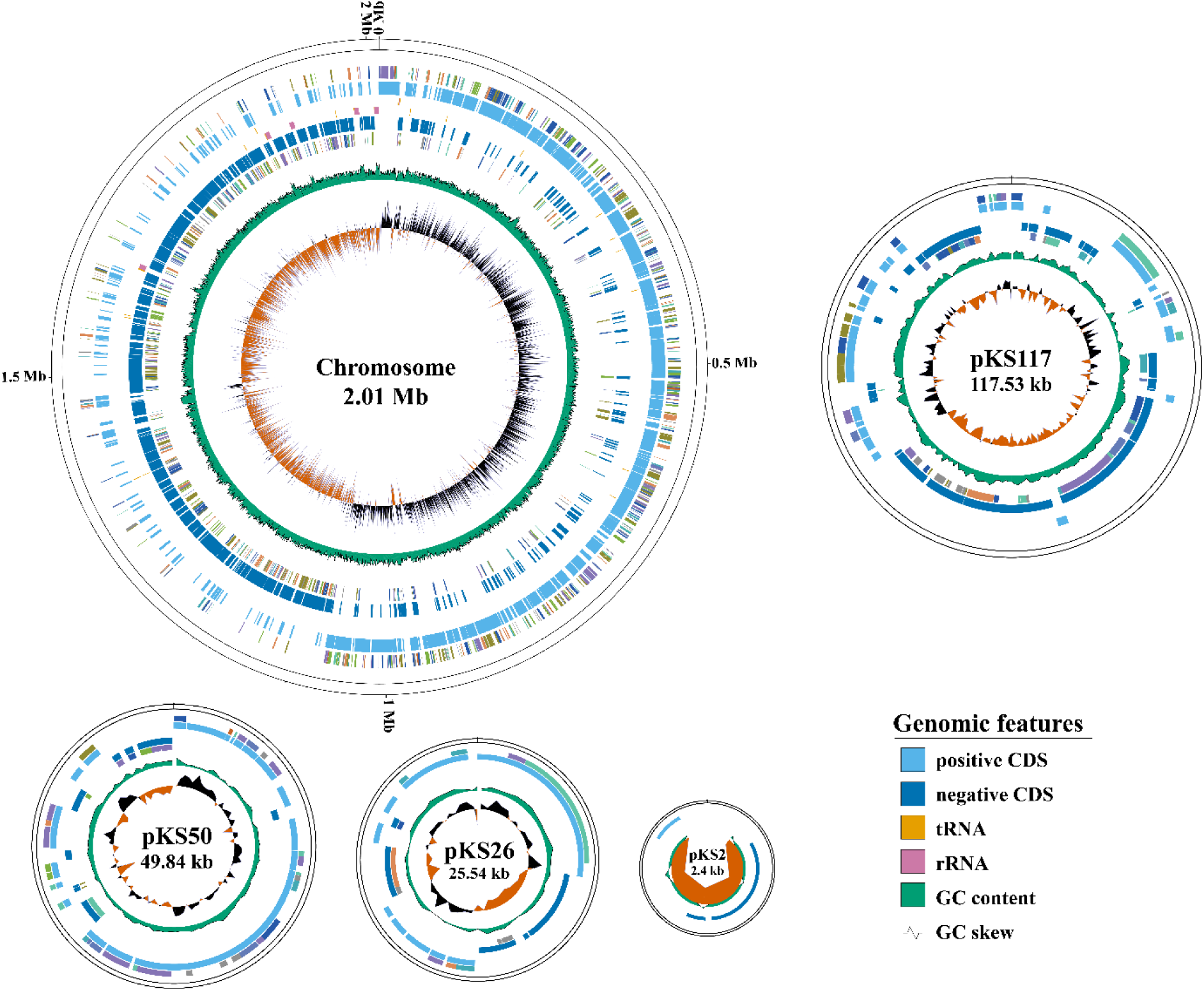
Circular genome maps of *L. garvieae* KS1546. The innermost ring shows the GC skew with black color indicating G-rich and orange C-rich. Coding sequences (CDS) on the leading strand are in light blue, reverse strand CDSs (dark blue). Figure was generated using GenoVi ^74^.

**Table 1.** Genome features of *L. garvieae* KS1546.

| Attribute | pKS2 | pKS26 | pKS50 | pKS117 | Chromosome |
| --- | --- | --- | --- | --- | --- |
| Size (bp) | 2 417 | 25 536 | 49 839 | 117 533 | 2 013 339 |
| GC content (%) | 33 | 35 | 32 | 34 | 38 |
| Genes | 5 | 30 | 61 | 130 | 2101 |
| CDSs | 4 | 30 | 60 | 127 | 1 988 |
| tRNAs | 0 | 0 | 0 | 0 | 60 |
| rRNAs | 0 | 0 | 0 | 0 | 16 |
| tmRNAs | 0 | 0 | 0 | 0 | 1 |
| ncRNAs | 1 | 1 | 1 | 3 | 8 |
| regulatory ncRNAs | 0 | 0 | 0 | 0 | 20 |
| *oriC/oriVs* | Yes | Yes* | Yes | Yes | Yes |
| pseudogenes | 0 | 0 | 0 | 0 | 9 |

\* No *oriV* was predicted by software

The GarKS biosynthetic genes were found to be located on plasmid pKS50, a 50 kb plasmid primarily encoding genes with putative functions related to DNA modification, repair, recombination and mobilization/conjugation.

Search for antibiotic-resistance genes revealed three genes similar to *mdt*(A), *lsa*(D) and *arr*, which are potentially involved in resistance to erythromycin/azithromycin/tetracycline, lincosamide/streptogramin and rifampin, respectively. All three genes were chromosomally encoded and showed 99% identity (100% sequence coverage) with *mdt*(A) (GenBank accession CAA63510), 94% identity (100% sequence coverage) with *lsa*(D) (NCBI accession WP_165713684.1) and 49% identity (76% sequence coverage) with *arr* (WP_000237816.1). Most likely, these genes are part of the core genome of *L. garvieae* / *L. petauri* ^75^. Regarding *mdt(A)*, however, it should be noted that the gene in our strain contained mutations previously shown to be associated with a lack of functional antibiotic resistance in *L. garvieae* ^76^. No virulence factor genes were identified in the KS1546 genome following BLAST screening against the VFDB dataset, using the EFSA-recommended thresholds of >80% amino acid identity and >70% query coverage.

### Functional confirmation of GarKS biosynthetic genes

The function of the GarKS biosynthetic genes have been inferred from homology and similarities with the genetic organization of the AurA70 cluster ^7,21^. Similar to AurA70, the biosynthesis of GarKS appears to involve three transcriptional units encoding the structural bacteriocin peptides (*gakABC*), putative immunity and regulatory proteins (*gakIR*), and a transporter protein (*gakT*) (Figure 2). However, the functions of these proteins have not been established experimentally. To confirm their predicted function, several constructs were made for heterologous expression of these proteins in *L. lactis* IL1403.

**Figure 2.**
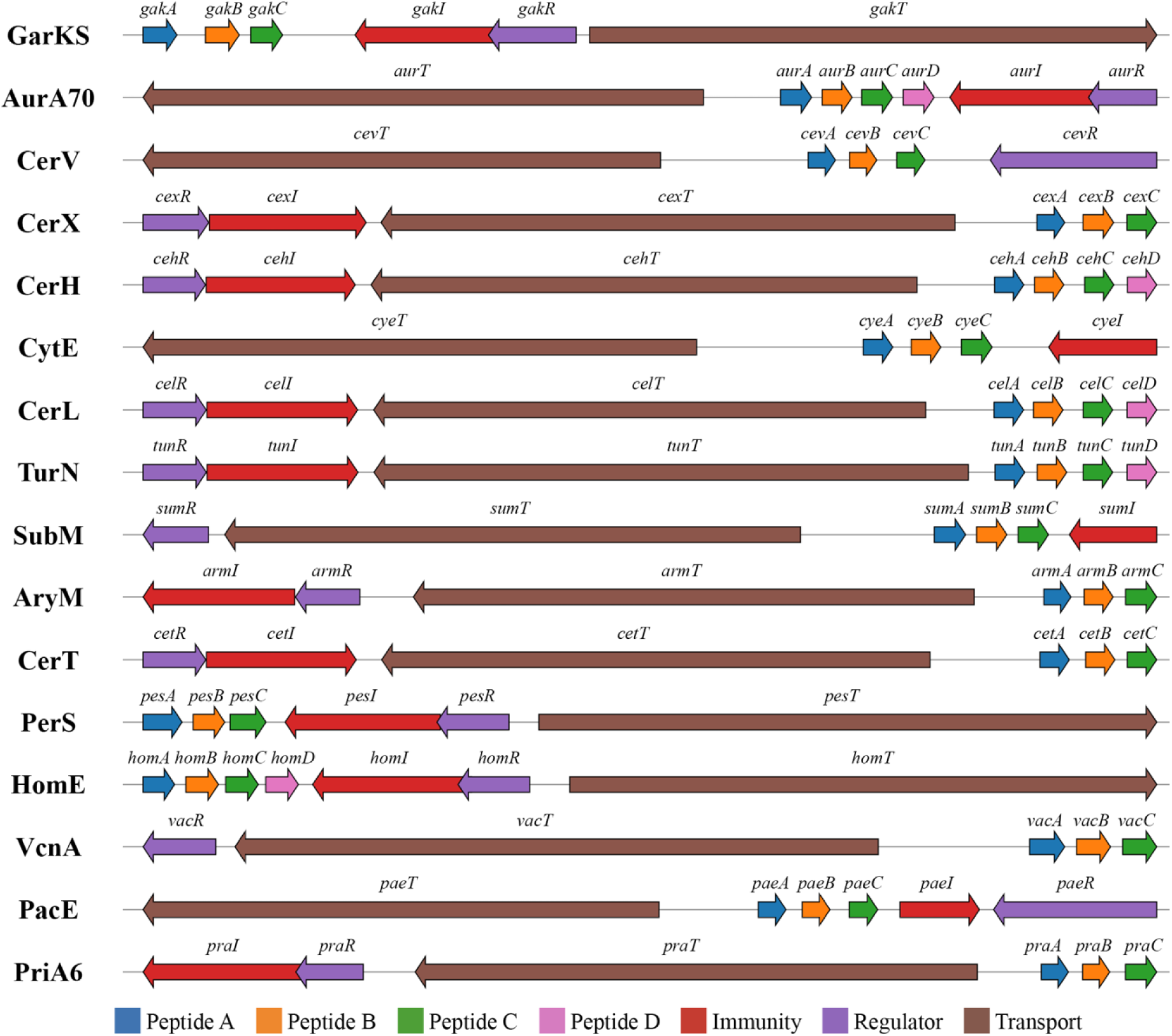
All multipeptide bacteriocin gene clusters. Figure was generated using the geneviewer v0.1.10 library for R v4.5.1.

To assess the role of *gakR*, the complete GarKS locus was cloned with and without *gakR* and the transformed culture was assayed for bacteriocin production. As shown in Table 2, the antimicrobial activity of culture supernatants from *L. lactis* IL1403 harboring the complete locus (*gakA2T*) was 480 BU/ml where one bacteriocin unit (BU) was defined as the lowest amount of bacteriocin needed to inhibit growth of *L. lactis* IL1403 by at least 90%. Comparably, the strain lacking *gakR* (*gakΔR*) showed a reduction in bacteriocin production by 6-fold to 80 BU/ml (Table 2).

**Table 2.**
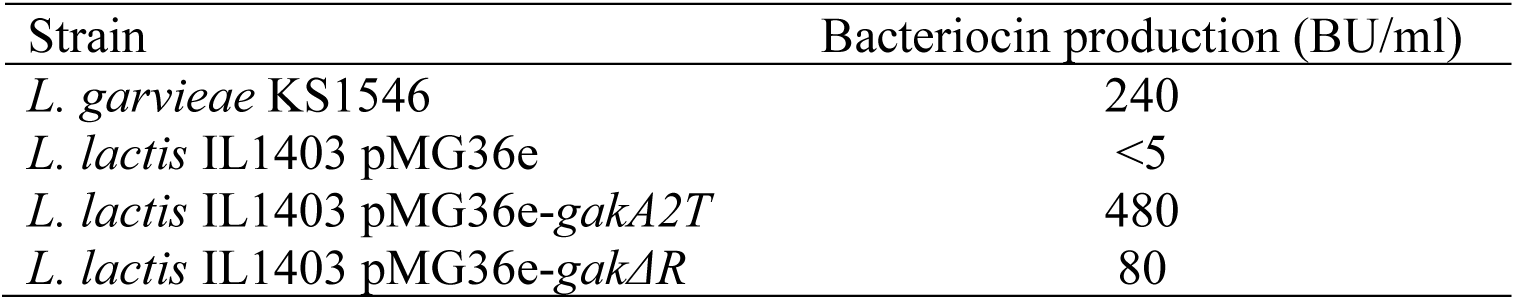
Comparative analysis of the GarKS production by the native producer and recombinant *L. lactis* IL1403.

The function of the predicted immunity protein was examined using *L. lactis* IL1403 transformed with a construct containing *gakI* (pMG36e-gakI), which was assayed for susceptibility to synthetic GarKS using a minimal inhibitory concentration (MIC) assay (Table 3). Expression of *gakI* in *L. lactis* IL1403 resulted in a 250-fold increase in the MIC for GarKS compared to a control strain harboring only pMG36e. We also tested whether GakI could cross-protect against the other bacteriocins in the same class, including new bacteriocins identified in this study (see below). An increase in MIC for cells harboring *gakI* compared to the control was also seen for the bacteriocins CerV, CerH, CerL, and SubM, but not AurA70, PerS or HomE.

**Table 3.** Antimicrobial activity of multipeptide bacteriocins against *L. lactis* IL1403 expressing the garvicin KS immunity protein *gakI*.

| Strain | MIC (nM) |  |  |  |  |  |  |  |
| --- | --- | --- | --- | --- | --- | --- | --- | --- |
|  | GarKS | AurA70 | CerV | CerH | CerL | SubM | PerS | HomE |
| *L. lactis* IL1403 pMG36e | 10 | 320 | 40 | 40 | 320 | 80 | 40 | 630 |
| *L. lactis* IL1403 pMG36e-*gakI* | 2500 | 320 | 1300 | 320 | 1300 | 1300 | 40 | 320 |

Heterologous expression of the *gak* locus without *gakT* (*gakΔT*) was also attempted, but no transformants could be obtained. We were also unable to obtain transformants with only the structural genes *gakABC*.

### GarKS susceptibility involves a PspC-domain containing protein

Bacteriocins are generally believed to disrupt bacterial cell membranes either by a receptor-mediated or receptor-independent mechanism. It is also generally thought that bacteriocins which require a receptor are far more potent than those acting by a receptor-independent mechanism ^77^. Given the high potency of some multipeptide bacteriocins, with MIC values below 5 nM, a receptor molecule could be involved in the mechanism of this family of bacteriocins. In a recent work, we found that in *L. monocytogenes*, a protein belonging to the phage shock protein response, influenced the susceptibility to GarKS, however, we could not identify any bacteriocins receptors ^48^. To further examine these aspects, we here attempted to isolate spontaneous GarKS resistant mutants of the susceptible strain *L. lactis* IL1403. Despite numerous efforts to isolate resistant cells, the highest level of resistance found was an 8-fold increase in MIC to 80 nM (Table 4). Whole-genome sequencing of the 8 most resistant isolates revealed mutations in *ythA*, encoding a phage shock protein C-domain-containing protein and a paralog to the protein involved in susceptibility in *L. monocytogenes* (UniProt accession Q9CED8); *ycbB*, a glycosyltransferase (Q9CIZ7); and *pstC*, a phosphate transport system permease protein (Q9CEW6). The highest decrease in susceptibility (increased MIC) was seen for two *ythA* mutants, both harboring a mutation near the end of the gene resulting in premature termination of translation and C-terminal truncation.

**Table 4.**
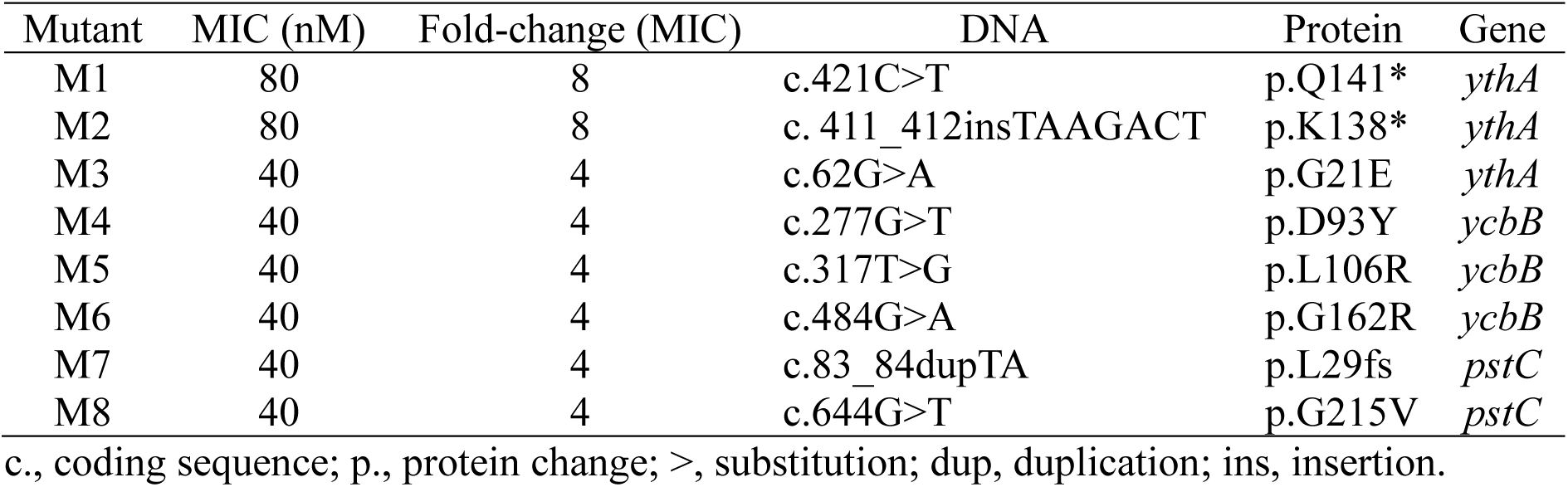
Mutations identified in spontaneous mutants of *L. lactis* IL1403 with reduced susceptibility to GarKS.

To further support the involvement of YthA in GarKS susceptibility, the *ythA* gene was introduced to mutant 1 (M1) for complementation and to *L. lactis* IL1403 as a second plasmid-encoded copy of the gene for overexpression. As shown in Table 5, the overexpression of *ythA* increased susceptibility to GarKS approximately 2.5-fold from 10 nM to 4 nM. The resistant mutant M1 became 4-fold more susceptible when complemented with a functional *ythA* gene, from 80 nM to 20 nM.

**Table 5.** GarKS susceptibility of *L. lactis* IL1403 and an *ythA* disruption mutant M1 expressing *ythA* from a recombinant plasmid.

| Strain | MIC<sub>90</sub> (nM) |
| --- | --- |
| *L. lactis* IL1403 pNZ8037 | 10 |
| *L. lactis* IL1403 pNZ8037-*ythA* | 4 |
| *L. lactis* M1 pNZ8037 | 80 |
| *L. lactis* M1 pNZ8037-*ythA* | 20 |

### Expanding the family of multipeptide bacteriocins

To our knowledge, only five multipeptide bacteriocins have been (partially) characterized to date, namely garvicin KS, aureocin A70 and the cereucins V, X and H ^7,21^. The limited number of biosynthetic gene clusters for multipeptide bacteriocins makes it difficult to identify conserved features such as regulatory mechanisms and accessory genes. Given the exponential rise of public sequence databases such as the NCBI nucleotide database and the recent release of Logan - an assembled version of the sequence read archive (SRA) - we sought to discover more multipeptide bacteriocins by genome mining ^70,78^. This led us to identify 11 putative new multipeptide bacteriocins (Table 6), hereafter named: cytocin E (CytE), cereucin L (CerL), thuricin N (TurN), subticin M (SubM), aryacin M (AryM), cereucin T (CerT), peromyscicin S (PerS), homicin E (HomE), vallicin A (VcnA), pacificin E (PacE), and priesticin A6 (PriA6).

**Table 6.**
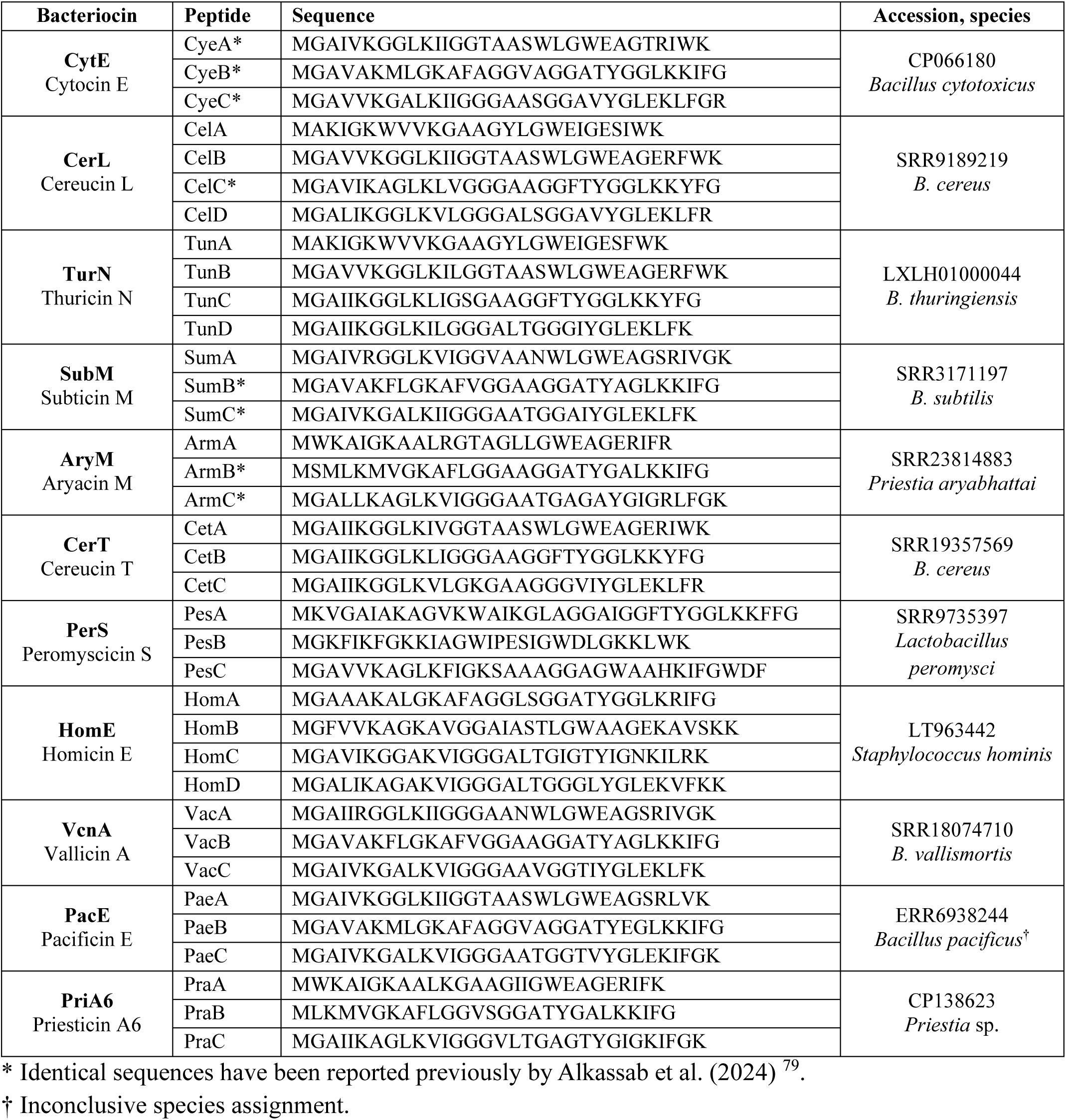
New multipeptide bacteriocins discovered by genome mining.

Most bacteriocins were encoded as part of seemingly complete biosynthetic clusters, which also contained genes putatively encoding a transcriptional regulator, an ABC-transporter, and an immunity-like gene (Figure 2). No putative regulator could be found near the genes encoding CytE, and no candidate for a gene encoding an immunity-like protein could be found for VcnA.

To confirm the antibacterial activity of the newly discovered bacteriocins, four of them, HomE, SubM, CerL and PerS, were obtained as synthetic peptides and assayed for antimicrobial activity against a panel of indicator bacteria using a minimal inhibitory concentration assay. Among all species tested, *L. lactis* was found to be susceptible to all four bacteriocins (Table 7). Activity was also observed for all four bacteriocins against *L. monocytogenes* and some isolates of *Enterococcus faecium*. The activity of the four bacteriocins against all indicators tested is shown in Supplementary Table S2.

**Table 7.**
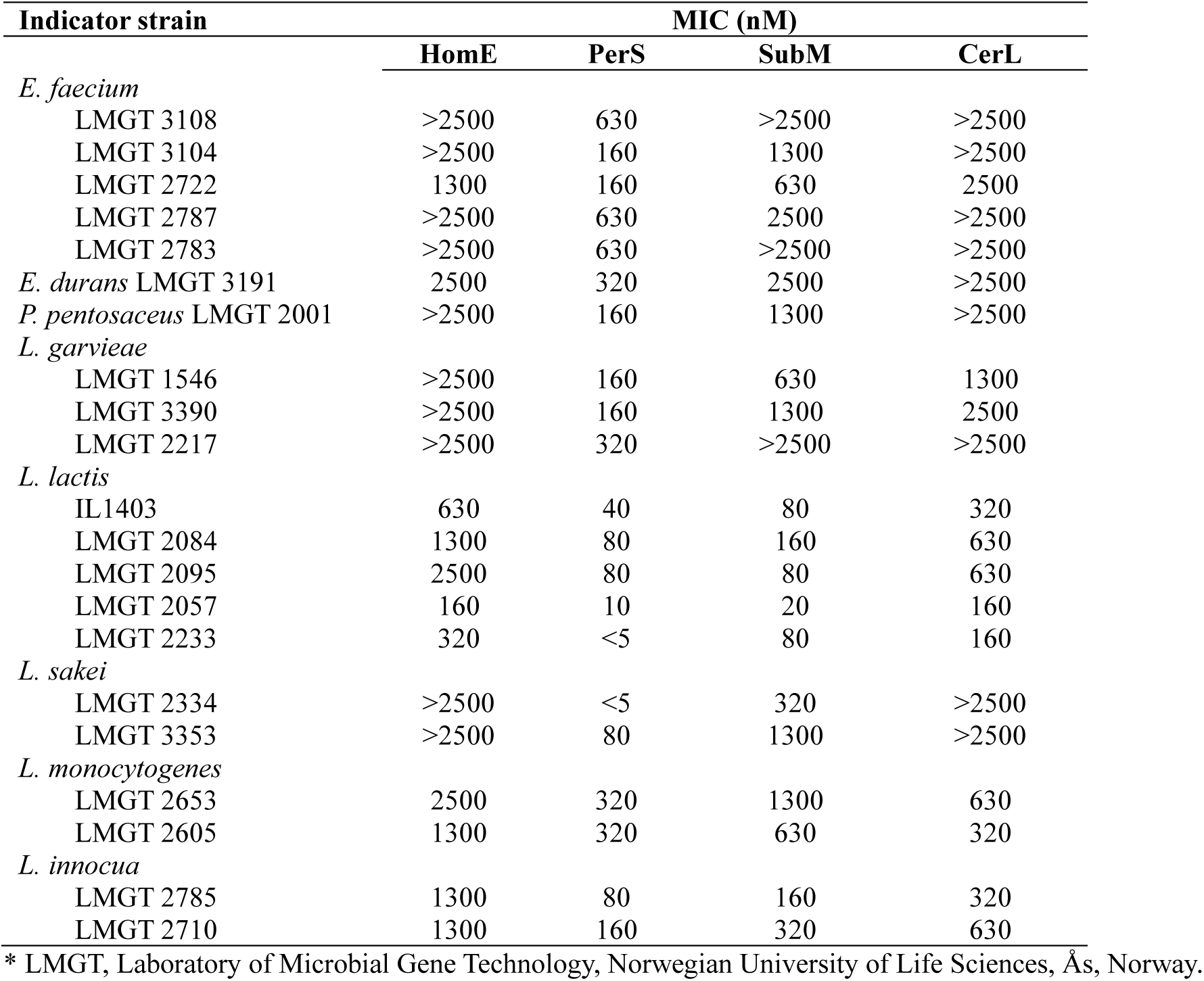
Antimicrobial activity (minimal inhibitory concentration) of new multipeptide bacteriocins against a panel of indicator species.

The multipeptide nature of the four bacteriocins was also assessed by assaying all combinations of the individual peptides constituting each bacteriocin (Table 8). For HomE, SubM and PerS, the lowest MIC was observed for the combination of all peptides, while for CerL, the combination of three and four peptides, namely ABD and ABCD, were equally potent (MIC of 160 nM). For SubM and PerS, the combination of all peptides had a MIC value 65- and 16-fold lower than the second most potent combination, respectively.

**Table 8.**
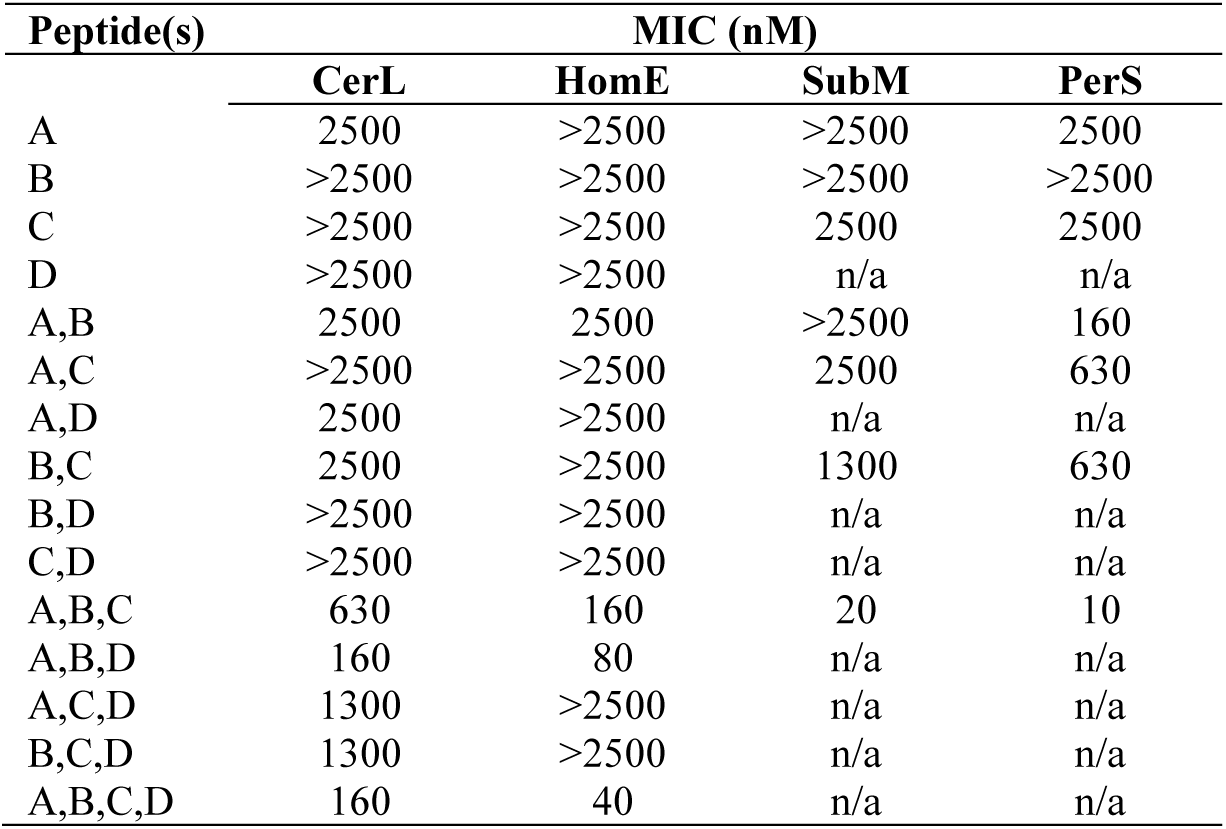
Activity of all peptide combinations of multipeptide bacteriocins against *L. lactis* IL1403 (minimal inhibitory concentration).

| Peptide(s) | MIC (nM) |  |  |  |
| --- | --- | --- | --- | --- |
|  | CerL | HomE | SubM | PerS |
| A | 2500 | >2500 | >2500 | 2500 |
| B | >2500 | >2500 | >2500 | >2500 |
| C | >2500 | >2500 | 2500 | 2500 |
| D | >2500 | >2500 | n/a | n/a |
| A,B | 2500 | 2500 | >2500 | 160 |
| A,C | >2500 | >2500 | 2500 | 630 |
| A,D | 2500 | >2500 | n/a | n/a |
| B,C | 2500 | >2500 | 1300 | 630 |
| B,D | >2500 | >2500 | n/a | n/a |
| C,D | >2500 | >2500 | n/a | n/a |
| A,B,C | 630 | 160 | 20 | 10 |
| A,B,D | 160 | 80 | n/a | n/a |
| A,C,D | 1300 | >2500 | n/a | n/a |
| B,C,D | 1300 | >2500 | n/a | n/a |
| A,B,C,D | 160 | 40 | n/a | n/a |

## Discussion

We have previously described the isolation of *L. garvieae* KS1546 from raw bovine milk which was shown to produce a multipeptide bacteriocin named GarKS. The bacteriocin was purified and shown to be active against a broad spectrum of bacteria, including food spoilage- and pathogenic bacteria ^7^. Furthermore, genetic engineering of the producer strain have been shown to increase the production of GarKS to over 1 g/L by batch fermentation using inexpensive growth media ^49^. Thus, GarKS shows high potential for medical and industrial applications, such as a component in wound dressings or as a biopreservative, either as pure peptide or as a crude fermentate, similar to the nisin-product Nisaplin. Prior to the use of a strain or its fermentates in such practical application, molecular level details of the strain safety profile as well as the bacteriocin biosynthesis, mode of action and potential resistance development is required. These aspects were investigated in the current work.

Our genome assembly of the strain resulted in a closed chromosome and four complete plasmids. Previous bioinformatic analysis of the garvicin KS producer identified the organism as *Lactococcus garvieae*, a species commonly associated with fish infections and occasionally reported in humans ^80–82^. However, comparative analyses with updated databases indicate that the genome is more accurately assigned to *Lactococcus petauri* (average nucleotide identity of 97.32%). This discrepancy reflects the close genetic relatedness of the two species, but more importantly the relatively recent description of *L. petauri* as a distinct taxon ^83^. This species was not represented in sequence databases prior to 2017, only after *L. garvieae* KS1546 was first described. This reclassification highlights the critical role of data availability when describing biological material such as bacterial isolates, as it allows for future bioinformatic analyses to revise and reconcile data with updated criteria. Notably, we identified the garvicin KS biosynthetic gene cluster on plasmid pKS50, a plasmid also encoding systems for both conjugal transfer and mobilization. This suggests a potential for horizontal transfer of the bacteriocin locus. However, we did not identify any other sequenced strains in the databases encoding GarKS.

The antibiotic resistance and virulence profile of the KS1546 strain is an important consideration when assessing the potential use of the strain or its fermentate in medical or industrial applications. By *in silico* analysis, we did not find that *L. garvieae / L. petauri* KS1546 encode any virulence factors, and the only potential antibiotic resistance determinants were found on the chromosome and are likely core genome members. These genes include, *lsa(D)*, a ribosome protection protein shown to confer resistance to lincosamides/streptogramins ^84^, *arr*, encoding an enzyme potentially inactivating rifampicin by ADP-ribosylation ^85^, and the putative drug exporter *mdt(A)* implicated in resistance to erythromycin and tetracycline. Regarding the latter, however, it should be noted that the *mdt(A)* variant encoded by KS1546 corresponds to the non-functional allele previously described in *L. garvieae*, which does not confer antibiotic resistance ^84^. Taken together, the complete genome sequence thus represents a valuable resource for future studies on the biology and biotechnological potential of *L. garvieae/L. petauri* KS1546.

The role of the GarKS biosynthetic genes can be predicted based on sequence homology and in this work we attempted to experimentally validate the functional predictions by heterologous expression. The protein encoded by *gakT* is highly similar to ATP-binding proteins containing both ATPase and permease domains of the multidrug resistance-like (*mdl*) family of ABC-transporters ^86^. We were not able to express a GarKS construct without the *gakT*, suggesting that expression of GarKS without GakT-mediated transport is toxic for the cells. Furthermore, GarR encodes a Cro/C1-type, helix-turn-helix domain-containing transcriptional regulator (InterPro domain IPR001387), a protein expected to play a role in regulating GarKS production ^7^. Indeed, our results showed that deletion of *gakR* reduces GarKS production six-fold, indicating that GakR functions as a positive regulator of GarKS biosynthesis. This is in contrast to the biosynthetic cluster of AurA70 in *S. aureus*, where a similar regulator, AurR, is thought to downregulate AurA70 production by binding to the promoter region of *aurABCD*^29^. Similarly, the cytolysin regulator CylR2 in *E. faecalis* binds as a homodimer to the promoter region to repress transcription of cytolysin genes ^30,31^. Such regulators commonly form homodimers and bind to inverted repeat sequences near promoters to regulate expression ^87^. Investigation of the upstream region of the *gakABC* operon revealed two 20-bp sequence regions that are inverted repeats spaced 2-bp apart (Figure 3). The spacing between the center of each sequence is 22-bp, corresponding to about two full helical turns of B-DNA. This spacing allows molecules bound to each repeat to face the same side of the DNA, consistent with binding of a homodimer ^88,89^. Furthermore, we have previously predicted the -35 and -10 elements of a possible *gakABC* promoter in the same region as the inverted repeat (Figure 3). Exploratory RNA-seq data from *L. garvieae / L. petauri* KS1546 is consistent with a promoter for *gakABC* at that position (data not shown). Structure prediction of the upstream region of *gakA* and two molecules of GakR using AlphaFold3 predicts binding of GakR to the inverted repeats as a homodimer (ipTM = 0.64, pTM = 0.68, Supplementary Figure S1). However, it is unclear why binding of GakR to this predicted site should result in transcriptional activation rather than repression due to the steric hindrance of RNAP and abortive initiation, as reported for AurR and CylR2 ^90^. Such divergent roles of homologous regulators underscore the diversity of bacteriocin regulation and suggest that regulatory strategies may be adapted to the ecological niches and physiological requirements of different producer strains. Understanding these mechanisms is important for the development of genetic engineering strategies to increase bacteriocin yield needed for industrial or therapeutic applications.

**Figure 3.**
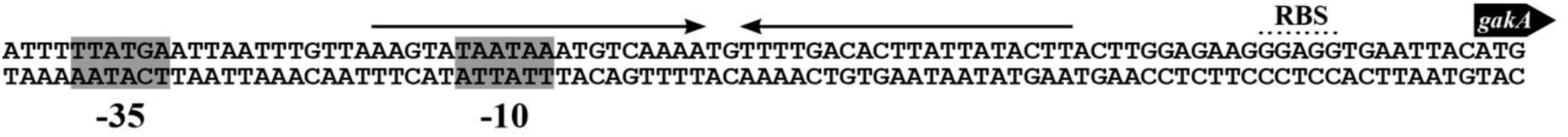
Predicted sequence elements in the upstream region of *gakA*. An inverted repeat indicated by arrows overlap with the -10 element of the predicted promoter.

Expression of the predicted immunity protein *gakI* in *L. lactis* IL1403 resulted in a 250-fold increase in the MIC_90_, clearly demonstrating its role in protection against the bacteriocin. The gene was also shown to confer protection against distantly related multipeptide bacteriocins such as CerV found in *Bacillus cereus* and SubM (*B. subtilis*), but not AurA70 from *S. aureus* or PerS from *Lactobacillus peromysci*. To explain this difference, we performed a multiple sequence alignment of all multipeptide bacteriocins, including the 11 new members identified in this work. This sequence alignment revealed three distinct clusters of sequence-related sequences, here named Group A, B and C (Figure 4), suggesting that multipeptide bacteriocins are essentially three-peptide bacteriocins with some having a duplicated gene. Indeed, one of the peptides in the four-peptide bacteriocin loci is often redundant and shows no or little synergy. For example, in our study, peptides A, B, and D for CerL show the same potency as peptides A, B, C and D, and for HomE the addition of peptide C increased potency only 2-fold (Table 8). It has also been shown that CehA is redundant for optimal activity of CerH ^7^. Among peptides belonging to group A, AurA and HomB, peptides from two bacteriocins that GakI does not protect against (AurA70 and HomE), are missing a conserved glutamic acid residue in a WExxE motif. The other bacteriocin PerS appears to be an outlier with no group A peptide and a group B peptide (PesB) quite divergent from the group (only 29.4% sequence identity with GakB at 85.3% coverage). These findings support the view that immunity specificity is determined by conserved structural motifs in the cognate bacteriocins, and that the mechanism of immunity is likely to involve direct interactions between the immunity protein and peptide(s).

**Figure 4.**
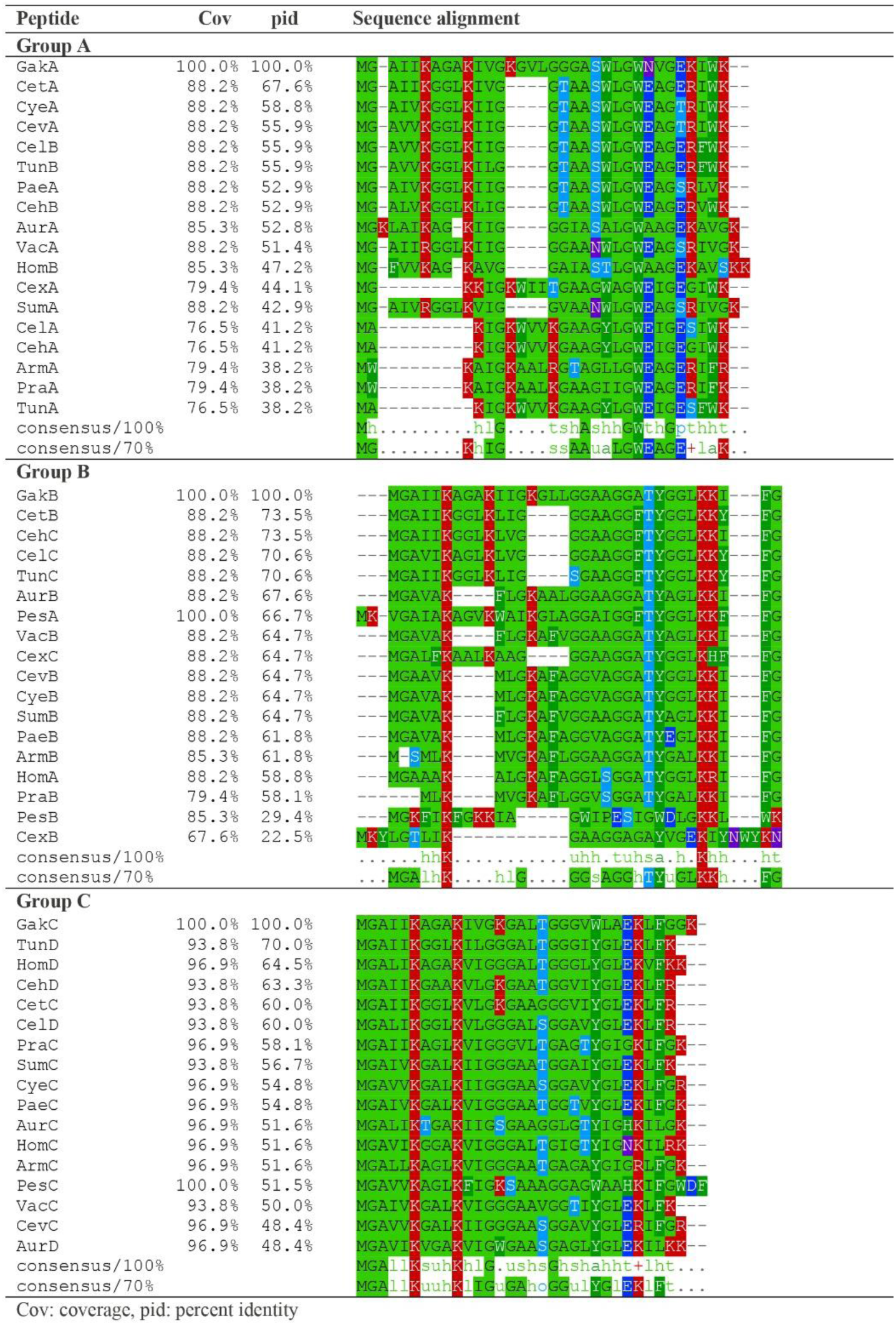
Multiple sequence alignment and clustering of multipeptide bacteriocins. Peptides were aligned using MAFFT (v7.511) ^91^ and visualized with MView ^92^. Cov: coverage, pid: percent identity.

As with *L. monocytogenes* ^48^, attempts to isolate *L. lactis* mutants with spontaneous resistance to GarKS did not yield highly resistant strains; the greatest reduction in susceptibility (8-fold) was seen for mutants M1 and M2 (Table 4) harboring mutations in *ythA*, a PspC-domain-containing protein. This result corroborates our previous finding of the PspC-domain protein Lmo2486 as the main GarKS susceptibility determinant in *L. monocytogenes,* showing that PspC-proteins modulate GarKS susceptibility across species. The *ythA* gene is the first member of an operon consisting of three genes *ythA*-*ythB*-*ythC*, predicted to encode a phage shock protein (Psp)-like system ^93^. The Psp response was first described in *E. coli* where it functions to protect the inner membrane and maintain energy homeostasis during envelope stress ^94^. In *L. lactis* one of the primary cell envelope stress response systems is the CesSR regulon, which is activated by cell envelope disturbances such as the cell wall inhibiting antimicrobial lactococcin 972 ^95^. Once activated, the transcriptional activator CesR binds to the promoter region of many genes which helps the cell restore cell envelope function and protect it from further damage ^93^. In support of this, mutants with constitutively active CesSR become resistant to lysozyme ^93^. Surprisingly, mutations in the *yth*-operon also causes lysozyme resistance and constitutive expression of the *ces*-operon ^93^. Due to these observations it has been speculated that YthA is involved in regulating the activation of CesSR, likely in a negative manner by protein-protein interactions to prevent over-induction ^93^. The decrease in susceptibility to GarKS in these mutants may therefore be a consequence of a constitutively activated CesSR rather than any direct interaction between GarKS and YthA, but this should be further investigated. We recently showed that the gene *lmo2486* is involved in GarKS susceptibility in *Listeria monocytogenes* EGD-e ^48^. The *ythA* gene from *L. lactis* and the *lmo2486* gene from *L. monocytogenes* likely encode functional homologs, both containing a PspC-domain. Similarly to *L. lactis*, *L. monocytogenes* with mutations in *lmo2486* show reduced susceptibility to GarKS, however, in this organism all reported mutants had introduced a premature stop codon in the *lmo2486* gene (K4NfsX). Thus, while the precise mechanism of cell targeting and killing by GarKS remains to be determined, the results here support the hypothesis of YthA/Lmo2486 as negative regulators of a stress response system which confers protection against GarKS across species.

In summary, the high potency, low resistance development and high level production capability of the multipeptide leaderless bacteriocin Garvicin KS, combined with the new collection of bacteriocins with different target spectra identified here, underlines the potential of this family of bacteriocins as promising candidates for different applications in biopreservation and infection treatment.

## Acknowledgments

The project was supported by The Research Council of Norway (Project: Bacpress **-** active packing technology to increase food shelf life, project number 341639) and funds from the Norwegian University of Life Sciences.

## Data availability

The complete annotated genome sequence of *L. garvieae* / *L. petauri* KS1546 has been submitted to NCBI with accession number [*in process*].

## Notes

### Competing Interest Statement

The authors have declared no competing interest.

