## Supporting information for "Novel insights into the biosynthesis and diversity of leaderless multipeptide bacteriocins"

**Table S1**. Primers used in this study.

| Primer | Sequence (5'-3') |
|---|---|
| A2T_SacI_F | CGTAATTCGAGCTCCACCTCTGCTGTTTTTC |
| A2T__HindIII_R | AGACTTTGCAAGCTTTTAATCCTGACTCATCAGATATTC |
| gakΔR_R | GCTTTATTTTGGAGGAAAGAATATG |
| gakΔRT_R | ATTCTTTCCTCCAAAATAAAGCCATTTCAATTAAATATAGGATACCAC |
| gakI_SacI_F | CGTAATTCGAGCTCGCAGTCACAAGAAATTGTGG |
| gakI_HindIII_R | CAGACTTTGCAAGCTTGTTCAAAAAGGTCGTAGCAC |
| gakABC_HindIII_R | AGACTTTGCAAGCTTGCAATATTACGTTTGTGGG |
| gakΔT_SacI_R | CAGACTTTGCAAGCTTCATTCCCGCAAATATTATATCG |
| ythA_XbaI_R | GCTCTAGATACTTTCCAGTGGCGAAATC |
| ythA_BamHI_F | TTGGATCCAGGAGGAAAATTATGTCTCAAAGACAATTAACAAAATC |

**Table S2**. Minimal inhibitory concentration of the multipeptide bacteriocins homicin E (HomE), peromyscicin S (PerS), subticin M (SubM), and cereucin L (CerL) against a panel of indicators.

| **Indicator strain** | **MIC (nM)** | | | |
|---|---|---|---|---|
| | **HomE** | **PerS** | **SubM** | **CerL** |
| *S. aureus* | | | | |
| LMGT 3310 | >2500 | 2500 | >2500 | >2500 |
| LMGT 3264 | >2500 | 2500 | >2500 | >2500 |
| LMGT 3260 | >2500 | 2500 | >2500 | >2500 |
| LMGT 3266 | >2500 | 2500 | >2500 | >2500 |
| LMGT 3305 | >2500 | 2500 | >2500 | >2500 |
| LMGT 3258 | >2500 | 2500 | >2500 | >2500 |
| LMGT 3289 | >2500 | 2500 | >2500 | >2500 |
| LMGT 3272 | >2500 | 2500 | >2500 | >2500 |
| *S. epidermidis* LMGT 3026 | >2500 | 2500 | >2500 | >2500 |
| *E. faecalis* | | | | |
| LMGT 3199 | >2500 | >2500 | >2500 | >2500 |
| LMGT 3330 | >2500 | 320 | 1300 | 2500 |
| LMGT 3359 | >2500 | >2500 | >2500 | >2500 |
| LMGT 3333 | >2500 | 630 | >2500 | >2500 |
| LMGT 3143 | >2500 | >2500 | >2500 | >2500 |
| LMGT 3351 | >2500 | >2500 | >2500 | >2500 |
| LMGT 3200 | >2500 | 630 | 2500 | >2500 |
| *E. faecium* | | | | |
| LMGT 3108 | >2500 | 630 | >2500 | >2500 |
| LMGT 3104 | >2500 | 160 | 1300 | >2500 |
| LMGT 2722 | 1300 | 160 | 630 | 2500 |
| LMGT 2787 | >2500 | 630 | 2500 | >2500 |
| LMGT 2783 | >2500 | 630 | >2500 | >2500 |
| *E. durans* LMGT 3191 | 2500 | 320 | 2500 | >2500 |
| *P. pentosaceus* LMGT 2001 | >2500 | 160 | 1300 | >2500 |
| *L. garvieae* | | | | |
| LMGT 1546 | >2500 | 160 | 630 | 1300 |
| LMGT 3390 | >2500 | 160 | 1300 | 2500 |
| LMGT 2217 | >2500 | 320 | >2500 | >2500 |
| *L. lactis* | | | | |
| IL1403 | 630 | 40 | 80 | 320 |
| LMGT 2084 | 1300 | 80 | 160 | 630 |
| LMGT 2095 | 2500 | 80 | 80 | 630 |
| LMGT 2057 | 160 | 10 | 20 | 160 |
| LMGT 2233 | 320 | <5 | 80 | 160 |
| *L. sakei* | | | | |
| LMGT 2334 | >2500 | <5 | 320 | >2500 |
| LMGT 3353 | >2500 | 80 | 1300 | >2500 |
| *B. subtilis* LMGT 3131 | >2500 | >2500 | >2500 | >2500 |
| *B. cereus* | | | | |
| LMGT 2805 | >2500 | >2500 | >2500 | >2500 |
| LMGT 2731 | >2500 | >2500 | >2500 | >2500 |
| LMGT 2711 | >2500 | 2500 | 2500 | 1300 |
| LMGT 2735 | >2500 | >2500 | >2500 | >2500 |
| *L. monocytogenes* | | | | |
| LMGT 319 | 2500 | 320 | 1300 | 630 |
| LMGT 2605 | 1300 | 320 | 630 | 320 |
| *L. innocua* | | | | |
| LMGT 2785 | 1300 | 80 | 160 | 320 |
| LMGT 2710 | 1300 | 160 | 320 | 630 |
| *S. hominis* LMGT 3129 | >2500 | 630 | >2500 | >2500 |
| *Lb. johnsonii* LMGT 3086 | >2500 | 160 | 2500 | >2500 |
| *Lb. delbrueckii* LMGT 3034 | >2500 | 320 | 2500 | >2500 |

* LMGT, Laboratory of Microbial Gene Technology, Norwegian University of Life Sciences, Ås, Norway.

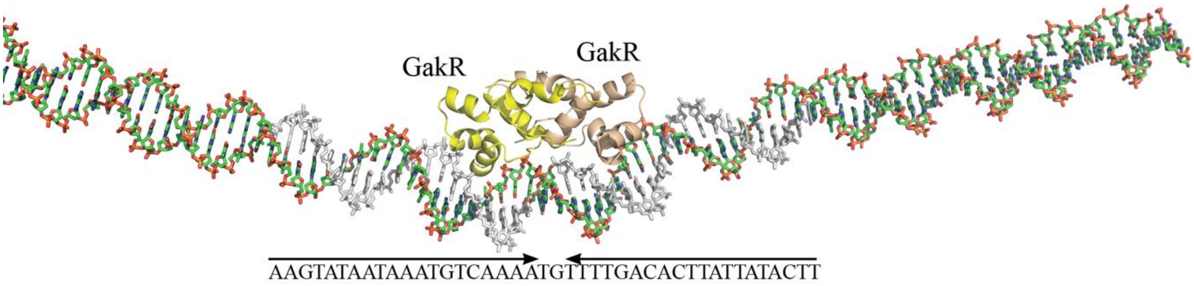


**Figure S1**. AlphaFold3 structure prediction of two GakR proteins and the DNA region immediately upstream of the *gakA* gene.
